# Stem cells in *Nanomia bijuga* (Siphonophora), a colonial animal with localized growth zones

**DOI:** 10.1101/001685

**Authors:** Stefan Siebert, Freya E. Goetz, Samuel H. Church, Pathikrit Bhattacharyya, Felipe Zapata, Steven H.D. Haddock, Casey W. Dunn

## Abstract

**Background:** Siphonophores (Hydrozoa) have unparalleled colony-level complexity, precision of colony organization, and functional specialization between zooids (i.e., the units that make up colonies) Previous work has shown that, unlike other colonial animals, most growth in siphonophores is restricted to one or two well-defined growth zones that are the sites of both elongation and zooid budding. It remained unknown, however, how this unique colony growth and development is realized at the cellular level.

**Results:** To understand the colony-level growth and development of siphonophores at the cellular level, we characterize the distribution of proliferating cells and interstitial stem cells (i-cells) in the siphonophore *Nanomia bijuga*. Within the colony we find that i-cells are present at the tip of the horn, the structure within the growth zone that gives rise to new zooids. They persist in the youngest zooid buds, but as each zooid matures i-cells become progressively restricted to specific regions within the zooids until they are mostly absent from the oldest zooids. I-cell marker-gene expression remained in gametogenic regions. I-cells are not found in the stem between maturing zooids. Domains of high cell proliferation include regions where i-cells can be found, but also include some areas without i-cells such as the stem within the growth zones. Cell proliferation in regions devoid of marker gene expression indicates the presence of mitotically active epithelial cell lineages and, potentially, progenitor cell populations.

**Conclusions:** Restriction of stem cells to particular regions in the colony may play a major role in facilitating the precision of siphonophore growth, and also lead to a reduced developmental plasticity in other, typically older, parts of the colony. This helps explain why siphonophore colonies have such precise colony-level organization.

## Background

Colonial animals provide a unique opportunity to investigate general questions about the evolution of development and to better understand development beyond embryogenesis [1–3]. Animal colonies arise when asexual reproduction is not followed by physical separation [4]. This results in many genetically identical multicellular bodies, known as “zooids”, that are attached and physiologically integrated. Colonial species are found in many animal clades, including ascidians, bryozoans, and many cnidarians [3]. The life cycles of colonial animals require multiple developmental processes - the embryological development of the zooid that founds the colony, the asexual development of subsequent zooids, and the colony-level development that regulates larger-scale colony formation including zooid placement [3].

Among colonial animals, siphonophores have both the highest degree of functional specialization between zooids and the most precise and complex colony-level organization [3]. In contrast to their benthic relatives, siphonophores have acquired a pelagic lifestyle and their zooids are arranged in very intricate repeating patterns along a linear stem (Figure 1). Each siphonophore colony has one or two main growth zones (depending on the species) where stem elongation takes place and new zooids arise by budding [5]. The localization of budding to such restricted zones and the consistency of budding within these zones results in very precise colony-level organization: in contrast to most other colonial animals, the zooids of a siphonophore are arranged in highly regular patterns that are are consistent between colonies of the same species. This budding process has been described at a gross scale for several species [6–8]. The youngest zooids are closest to the growth zone and the oldest are furthest from it, providing complete ontogenetic sequences of zooid development within a colony. This greatly facilitates developmental studies. Nothing is known, however, about the cellular dynamics of siphonophore colony growth. It is not known which regions have actively dividing cells, and the distributions of stem cells have never been described in siphonophores. This means that their potential role in zooid budding and colony elongation remain unknown.

**Figure 1.**
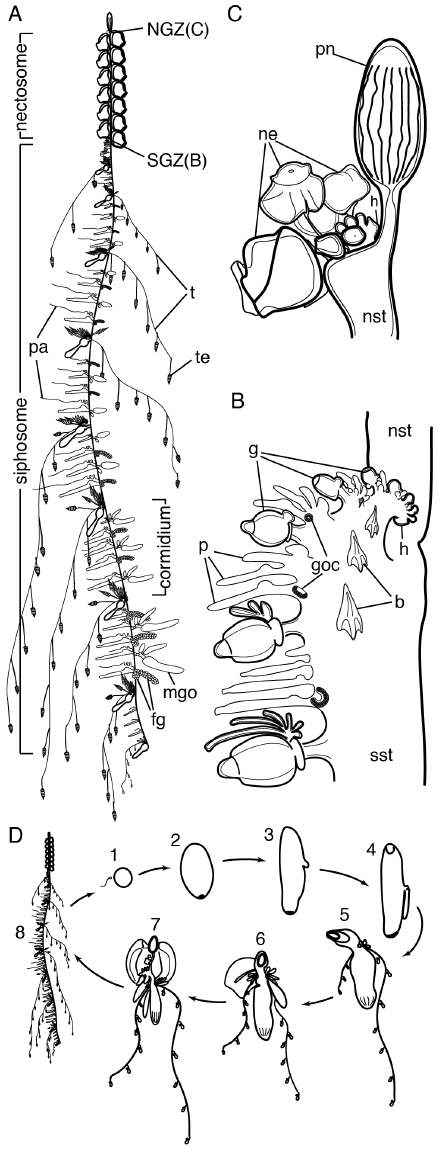
Schematic of *Nanomia bijuga*. Anterior [50] is towards the top of the illustrations. (A) Colony stage of the life cycle. For clarity reasons, protective bracts were not pictured and gonodendra of only one sex are shown per palpon in older parts of the colony. Approximate length of the illustrated colony was 15cm. The side of zooid attachment within the siphosome is defined as the ventral side of the stem [50]. (B) Siphosomal growth zone and anterior part of the siphosome. Sites of gonodendra formation (goc) are located at the bases of young gonodendra (shown here only for the most posterior palpon in each cormidium). Gonodendra mature in older cormidia further to the posterior (see A). (C) Nectosomal growth zone with the gas filled floating organ, the pneumatophore, at the top. (D) Lifecycle of *Nanomia bijuga*. 1. Egg and sperm. 2. 1.5 day old planula. 3. Two day old planula with larval tentacle bud. 4. 2.5 day old planula with forming pneumatophore and developing larval tentacle. The mouth opening of the protozooid is at the bottom. 5. One week old siphonula with pneumatophore and two larval tentacles bearing larval tentilla. 6. 20 day old siphonula with larval bract, and zooids developing on the ventral side of the protozooid indicating that growth zones have been established. 7. Young colony with first functional nectophore and zooids present along the elongating body of the protozooid. The elongating body of the protozooid corresponds to the future stem of the polygastric stage. 8. Mature colony - polygastric stage with multiple gastrozooids. Original figure was adapted from [27]. b: bract; fg: female gonodendron; g: gastrozooid; goc: gonodendral i-cell cluster; h: horn; mgo: male gonophore; ne: young nectophores; NGZ: nectosomal growth zone; nst: nectosomal stem; p: palpon; pa: palpacle, pn: pneumatophore; SGZ: siphosomal growth zone, sst: siphosomal stem; t: tentacle; te: tentillum. A-C modified from [51]. D modified from [52].

Stem cells were first described in hydrozoans [9], the clade that includes siphonophores, where they are referred to as interstitial stem cells (i-cells) since they are located within interstices between epithelial cells. Among colonial hydrozoans, i-cells have been studied in the greatest detail in *Hydractinia echinata* and *Clytia hemisphaerica* [10–12]. In *Hydractinia*, they give rise to all cell types (including epithelial cells). These i-cells are found throughout the colony and facilitate growth at different sites [2, 10], depending on environmental conditions. Hydrozoan i-cells have a distinct round or spindle shape, an enlarged nucleus, and chromatin that is less dense than that of other cells [13], which makes them conspicuous in micrographs. They also have characteristic gene expression profiles [12, 14–16]. Since siphonophores can only add new zooids within well-defined locations unlike most other colonial hydrozoans, it is important to know if stem cells are also restricted to particular regions, or widely distributed as in these other species. Spatial restriction of stem cells could have a mechanistic role in restricting zooid addition in siphonophores, enabling their precise and complex growth.

Here we identify regions of cell proliferation and describe the expression of i-cell marker genes in the siphonophore *Nanomia bijuga* (Figure 1). These observations allow us to answer fundamental questions about colony-level development in siphonophores.

## Methods

### Collection of Nanomia Bijuga Specimens

*Nanomia bijuga* specimens were collected from the floating dock in front of Friday Harbor Labs (FHL), San Juan Island, WA (12–19 June 2011), and in Monterey Bay, CA, and adjacent waters. Monterey Bay specimens were collected on 29 Sep 2012 via blue-water diving from a depth of 10–20 m and on 28 Sep to 03 Oct 2012 by ROV Doc Ricketts (R/V Western Flyer) at depths ranging from 348 – 465m. After collection, specimens were kept in filtered seawater (FSW) overnight at 8°C in the dark. Specimens for EdU labeling were collected on 19 Mar 2014 in Monterey Bay by ROV Ventana (R/V Rachel Carson) at depth ranging from 154–377m and on 23 May 2014 by ROV Doc Ricketts (R/V Western Flyer) at a depth of 300m. No ethical approval was needed as *Nanomia bijuga* is not subject to any animal care regulations.

### Identification and Amplification of Interstitial Stem Cell/Germ Cell Marker Genes

We used tblastx to identify *Nanomia bijuga* homologs for *piwi*, *nanos-1*, *nano-2*, *vasa-1* and *PL10* in a *Nanomia bijuga* transcriptome reference using available sequence information from *Clytia hemisphaerica*, *Podocoryne carnea* and *Hydra vulgaris*. Sequences for the *Nanomia bijuga* orthologs have been submitted to Genbank (Accession Nos. KF790888-790893). The transcriptome reference was in parts based on raw reads available at the Short Read Archive, Accession No. SRR871527 [17].

### Sequence Alignments and Phylogenetic Analysis

For each gene, a subset of significant RefSeq blast hits that matched the sampling in Kerner et al. [18] was used for phylogenetic analyses. We used MUSCLE v3.8.31 [19] to generate multiple sequence alignments for each gene separately, except for PL10 and vasa that were combined into a single matrix because they are sister gene families [18]. RAxML v7.5.7 [20] was used for phylogenetic analysis with the WAG model of amino acid substitution and the Γ model of rate heterogeneity. We used the non-parametric bootstrap [21] with 500 replicates for each matrix to assess support on each gene tree. The source code for the phylogenetic analyses, as well as the input fasta sequence files for all considered sequences and the output trees in newick format, are available as a git repository at https://bitbucket.org/caseywdunn/siebert-etal. Complete program settings can be found within these files.

### Whole Mount RNA in Situ Hybridization

*In situ* hybridization was performed on both Friday Harbor and ROV collected specimens and yielded consistent expression patterns. Four colonies per gene in four independent rounds of *in situ* hybridization were analyzed. ROV specimens are presented in the figures since their gonodendra were more mature. Specimens were transferred into a Petri dish coated with Sylgard 184 (Dow Corning Corporation) and relaxed by adding isotonic 7.5 % MgCl_2_·6H_2_O in Milli-Q water at a ratio of approximately ⅓ MgCl_2_ and ⅔ FSW. After pinning them out in a stretched position using insect pins (Austerlitz Insect Pins, 0.2mm, Fine science tools) they were fixed in 0.5% glutaraldehyde/4% paraformaldehyde (PFA) in FSW for two minutes and incubated in 4% PFA in FSW overnight at 4°C. Mature nectophores and bracts tend to get detached when handling specimens in the dish and were therefore not accessible for analysis in all cases. Specimens were then washed three times in PTw (phosphate buffer saline and 0.1% Tween). Dehydration was performed using EtOH with 15 min washes in 25% EtOH/PTw, 50% EtOH/PTw, 75% EtOH/Milli-Q water, 2x 100% EtOH and then transferred to MetOH and stored at −20°C. Use of EtOH for dehydration was empirically found to minimized tissue sloughing, detachment of endoderm from ectoderm.

Dig-labeled probes were generated using Megascript T7/SP6 kits (Life Technologies). Probe lengths in base pairs were as follows: *nanos-1*: 800; *nanos-2*: 954; *PL10*: 1,233, *piwi*: 1, 389; *vasa-1*: 1,381. Primer used for probe generation were as follows: *nanos-1_F* GAA CAC TCG CTA GTT GCT GTG, *nanos-1_R* TCT ATC GGT TTT AAC TTT TGG TG; *nanos-2_F* AGT AGT GGG AGC AGC CAA TG, *nanos-2_R* AAC CGT TGG TGG ATT GAT TC; PL10_F ACT GCT GCA TTT TTG GTT CC, *PL10_R* TGC CTG TTG CTG GTT GTA TG; *piwi_F* CAT GCT GTG TGC TGA TGT TG, *piwi_R* GCA AAG GCC TCT TTG AAT TG; *vasa-1_F* TTC CGG ACT ATT GCT CAA GG, *vasa-1_R* GAT CCC AGC CAT CAT CATT C. Working concentration of mRNA probes were 1ng/ml. *In situ* hybridizations were performed according to the protocol described by Genikhovich and Technau [22] with few deviations. Starting at step #27, the specimens were incubated in MAB instead of PTw. The blocking buffer composition was MAB with 1% BSA and 25% sheep serum. Anti-Digoxigenin-AP, Fab fragments (Cat.No.11093274910, Roche Diagnostics) were used in 1:2000 dilution in blocking buffer. After antibody binding the specimens were washed in MAB instead of PBT. Once the NBT/BCIP development was stopped with water, the samples were stored overnight in 100% ethanol followed by storage in PBS. Samples were stable in PBS for many weeks provided that the medium was exchanged regularly to prevent bacterial growth. Photodocumentation was performed using Canon MP-E 65mm Macro lens or using stereomicroscope Leica S8APO. In Figure 2A and Additional file S2C, a stacking strategy (focal montaging) was applied to increase depth of field. Four photographs with different focal planes were merged using function “auto blend layers” in Adobe Photoshop CS 5.5. After all photo documentation was completed, specimens were stored in 4% PFA/PBS and the integrity of the signal has remained stable. This is a preferable long-term storage because the tissue structure is preserved. Large specimens were difficult to mount because of size of tissue fragments. When trying to permanently mount tissue in Euparal (BioQuip Products, Inc) the mounting procedure caused tissue damage and over time strong unspecific staining occurred despite several washes in water after stopping the staining reaction.

**Figure 2.**
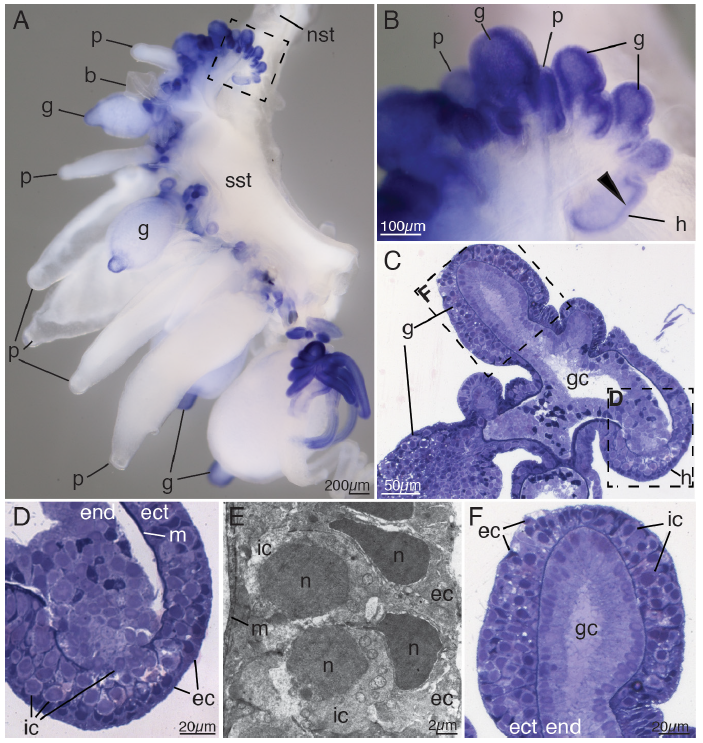
Siphosomal growth zone. (A) Anterior part of the siphosome, stained blue for *vasa-1* transcript. Lateral view. Anterior is up, ventral to the left. (B) Close-up of growth zone region boxed region in A. Marker gene expression within the horn (h). (C) Semi-thin longitudinal section of the tip of the siphosomal growth zone stained with toluidine blue. (D) Siphosomal horn, close-up of box in C. (E) Transmission electron micrograph of the ectoderm of the siphosomal horn. Interstitial cells reside in between epithelial muscle cells of the ectoderm. (F) Tip of youngest gastrozooid, close-up of box in C. b: bract; ec: epithelial cell; ect: ectoderm; end: endoderm; g: gastrozooid; gc: gastric cavity; h: horn of the growth zone; ic: interstitial cell; m: mesoglea; n: nucleus; nst: nectosomal stem; p: palpon; sst: siphosomal stem.

### Thick and Ultrathin Sectioning for Transmission Electron Microscopy

Specimens fixed as described above were washed with PBS five times for 15 min each and afterwards stored at 4°C in the presence of sodium azide ([1ng/ml]). Specimens were postfixed in 2% glutaraldehyde, 4% paraformaldehyde, 100mM sucrose and 100 mM sodium cacodylate buffer (SCB) overnight at room temperature. After three washes in 100 mM sucrose, 100mM SCB for 15 min each samples were postfixed in 1% OsO4, 100mM sucrose, 100mM SCB overnight at room temperature. Tissue was processed for resin embedding according to the manufacturer’s instructions (Low viscosity embedding Kit, Cat. 14300, Electron Microscopy Sciences). All washes and incubations were conducted at slow agitation on a rocker table. Thick sections (0.5–0.750μm) were prepared using glass knives, dried and counterstained for 30 seconds in toluidine blue (0.1%) in sodium borate (1%) buffer. Ultra thin sections were prepared using a diamond knife. Transmission electron microscopy (TEM) images were acquired on Phillips 410 Transmission Electron microscope. A representative set of thick sections was deposited at the Museum of Comparative Zoology, Harvard University (catalog numbers IZ50112-50113).

### Click-iT Cell Proliferation Assay

After collection, specimens were kept at 5–7°C overnight or up to 2d in the dark. Each specimen was truncated to a colony length of approximately 8cm in relaxed state by surgical removal of posterior parts of the siphosome. This was done to ensure comparable amounts of tissue in different incubations. Colonies (C1–C6) were incubated in 50ml volume per individual at click-iT® EdU concentrations of 100μM (5 specimens, C1–C5) and 20μM (1 specimen, C6) in FSW for 5h at a temperature of 5–7°C. Both concentrations yielded comparable results. Specimens were transferred into a Petri dish coated with Sylgard 184 and fixed as described above for *in situ* hybridization specimens. Dehydration was performed using EtOH with 15 min washes in 25% EtOH/PTw, 50% EtOH/PTw and 2x 75% EtOH/Milli-Q water and specimens were stored at −20° Celsius. To compare cell proliferation in different regions of the colony the specimens were dissected prior to the click-iT reactions. The nectosomal and the siphosomal growth zones including adjacent stem regions and up to three siphosomal fragments (SF1, SF2, SF3) with fully-grown zooids attached to the stem were transferred into wells of a 24 well plate (Costar 3524, Corning Incorporated). Siphosomal fragments (SF1, SF2, SF3) were taken at distances of approximately 1.5cm, 3cm and 4.5cm in posterior direction from the siphosomal horn and included at least one mature gastrozooid. The stem length in these tissue samples varied in between 2.5mm and 1cm. The tissue was rehydrated and permeabilized at room temperature using 10min washes in 50% EtOH/PBS, 25% EtOH/PBS, 2x PBS, 2x 3% BSA in PBS, 0.5%Triton X in PBS (20min) and 2x 3% BSA in PBS. The click-iT reaction was performed according to the manufacturer’s instructions (Click-iT® EdU Alexa Fluor® 594 Imaging Kit, C10339, Life Technologies). Before mounting the tissue was counterstained with DAPI (D1306, Life Technologies) solution at a concentration of 2ng/μl. The tissue was mounted in Vectashield (H–1000, Vector laboratories) and analyzed on a Zeiss LSM 510 Meta Confocal Laser Scanning Microscope. The overview shot presented in Figure 7A was generated manually in Adobe Photoshop CS6 by merging eight individual shots, which were taken consecutively using an identical focal plane. Comparisons between zooids of different developmental stages were made within one specimen when possible. The fixation and mounting procedure however rendered particular tissues inaccessible for confocal analyses in some cases. Photographs for presentation purposes had therefore be acquired across colonies on few occasions. Colonies analyzed are indicated in the figure legend. Photographs shown in Figure 7 summarize observations made across all six analyzed specimens. A quantitative analysis of cell division was however not accessible at this stage. Images were taken of tissue of colony C1 (Figure 7A–C,G,I–J,L–O,Q–R), colony C2 (Figure 7F,K), colony C3 (Figure 7P,S–W) and colony C6 (Figure 7D–E,H), respectively

## Results

We identified i-cells in siphonophores by examining the expression of five genes frequently used to identify i-cells in other hydrozoans [2, 12, 14–16] – *nanos-1*, *nanos-2*, *PL10*, *piwi* and *vasa-1*. Broadly-sampled phylogenetic analyses indicate that the sequences we identified in *Nanomia bijuga* are orthologs of these genes (Additional file 1A–C). Negative controls with sense probes were performed for *in situ* hybridizations of all genes in all zooids, and none were positive (Additional files 2–6). We confirmed the presence of cells with i-cell morphology in several regions that showed high expression of i-cell genes.

### I-Cells are Present in the Horn of the Siphosomal Growth Zone

The siphosomal growth zone produces most zooids in *Nanomia bijuga* (Figure 1A,B). The general structure of the *N. bijuga* siphosomal growth zone, as well as its budding process, has previously been described [7]. The zooids are arranged in repeating groups, known as cormidia. The budding sequence that produces cormidia and the zooid arrangement within them are highly organized (Figure 1A, [7]). The siphosomal growth zone has a protrusion at its anterior end - the horn (labeled h in Figure 1B). Pro-buds form at the tip of the horn and then subdivide into zooid buds as they mature and are carried to the posterior. These buds give rise to five different zooid types - gastrozooids (feeding polyps), palpons (polyps with function in circulation, defense, sensing and digestion), bracts (defense), and female and male gonophores (gamete production) [5].

All examined genes were found to be expressed at the tip of the siphosomal horn and in all buds and young zooids within the siphosomal growth zone (Figure 2A,B, Additional files 2C, 3C, 4C, 5C, 6C). Semi-thin sections and TEM analysis confirmed the presence of two types of cells within the ectoderm of the siphosomal horn, epithelial cells and i-cells (Figure 2C–E). Within the horn, cells with i-cell morphology were found in the endoderm (Figure 2D). The mesoglea within the horn appeared discontinuous suggesting that there may be migratory activity of i-cells between ectoderm and endoderm (Figure 2D). In the endoderm of young zooid buds, however, no cells with i-cell morphology were observed. Their nuclei were located close to the mesoglea (Figure 2F). Both epithelial cells and i-cells were found in the ectoderm of young zooids (Figure 2C,F).

### I-Cells Become Spatially Restricted During Zooid Development and are Largely Absent from Mature Zooids

The distal portion of the pro-bud gives rise to the gastrozooid – the feeding zooid (Figures 1B, 2B, 3). Young gastrozooid buds had strong expression of all marker-genes (Figure 3A, Additional files 3C, 4C, 5C, 6C). The basigaster, a specialized region of nematocyst formation in siphonophores [5], was evident in young gastrozooid buds as a thickening of the proximal ectoderm (Figure 3B). In the course of basigaster development, expression of all examined genes, except *nanos-2* (Figure 3F), became restricted to deep basigaster ectoderm (Figure 3B–E, Additional files 4C-D, 5C, 5E, 6C, 6E-F) and then decreased until a signal was no longer detectable (Figure 3H, Additional files 4G, 5E-F, 6F-G). *nanos-2* expression persisted in the basigaster region of gastrozooids of all ontogenetic stages (Figure 3F, Additional file 3E,G–H). This finding was consistent with previous studies that indicated a *nanos-2* function in nematocyst formation [12, 23]. Within the basigaster *nanos-2* seemed to be colocalized to the same region as minicollagen (see [24]), which is known to be involved in capsule formation [25]. Though *vasa-1, PL10, nanos-1* and *piwi* transcripts were not detected in basigasters of mature gastrozooids (Figure 3H, Additional files 4G, 5F, 6F), undifferentiated cells were still found along the mesoglea (Figure 3G) indicating the presence of a determined progenitor cell population which gives rise to nematocytes but has lost interstitial stem-cell transcriptional signatures. Immature nematocysts were observed in the outer layers of the mature basigaster (Figure 3G). The gene *vasa-1* was expressed in the same regions of the young gastrozooids as *PL10, nanos-1* and *piwi*. In addition, it was expressed in both the ectoderm and endoderm of the tips of the young gastrozooids (Figures 2A, 3B–E).

**Figure 3.**
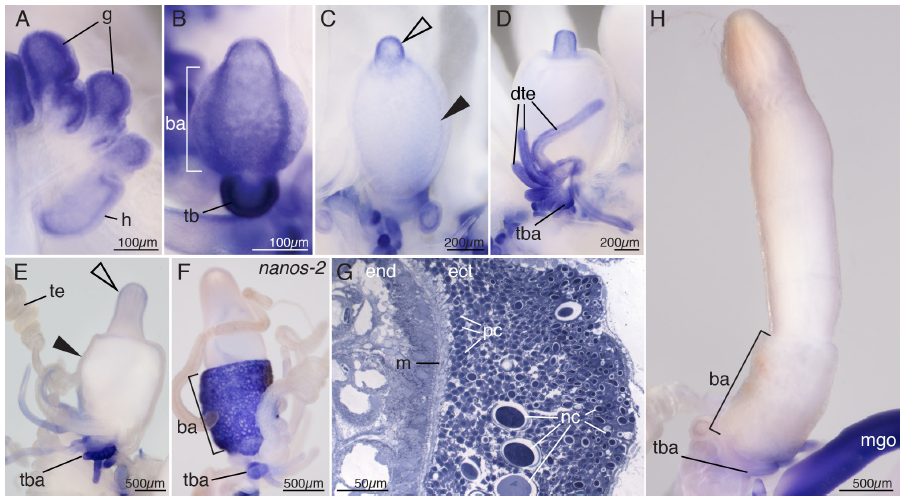
Gastrozooid development. (A-E, H) Ontogenetic series of gastrozooids, stained blue for *vasa-1* transcript. Distal is up. (A) Young gastrozooid buds close to the siphosomal horn. (B) Young gastrozooid with strong *vasa-1* expression in the developing tentacle bud. Within the basigaster region transcript was found predominantly in deeper tissue layers. Anterior view. (C) Slightly older gastrozooid with *vasa-1* expression in the gastrozooid tip (open arrowhead) and faint signal in the basigaster region (filled arrowhead). Posterior view. (D) Early stage of tentacle formation with developing tentilla branching off the tentacle. Anterior view. (E) *vasa-1* transcript disappears from maturing gastrozooid within the developing tip (empty arrowhead) and from the basigaster region (filled arrowhead) but remains present in tentacle bases and developing tentilla. Lateral view, anterior to the left. (F) *nanos-2* expression in the basigaster region and the tentacle base of a gastrozooid. Lateral view, anterior to the right. (G) Semi-thin longitudinal section of a mature gastrozooid basigaster, stained with toluidine blue. Undifferentiated cells (pc) are present along the mesoglea in ectodermal tissue. (H) Mature gastrozooid, stained blue for *vasa-1* transcript. *vasa-1* expression is not detectable in the body of the mature gastrozooid. Lateral view, anterior is to the right. ba: basigaster; dte: developing tentilla; ect: ectoderm; end: endoderm; g: gastrozooid; h: horn; m: mesoglea; mgo: male gonophore; nc: developing nematocysts; pc: putative nematocyte progenitor cells, tb: tentacle bud; tba: tentacle base; te: tentillum.

Each gastrozooid has a single tentacle attached at its base. The tentacle has side branches, known as tentilla, which bear packages of nematocysts at their termini (Figure 1A, [5]). All marker genes were expressed in the tentacle bases throughout all ontogenetic stages of gastrozooids (Figures 3B, 3D–F,H, Additional files 3H-I, 4G, 5F, 6G). The expression domains, however, differed between marker-genes. Whereas *nanos-2* expression was restricted to the very proximal end of the tentacle and very early tentilla buds (Figure 3F, Additional file 3H-I), signal for the other four genes persisted in developing tentilla as well (Figure 3D-E, Additional files 3E, 4G, 5F, 6G). Marker-genes were not expressed in the mature tentilla.

Anterior to each gastrozooid, a series of buds develop into palpons – zooids thought to have a function in circulation of gastrovascular content, digestion, defense and sensing (Figure 1B, [26]). Like gastrozooids, each palpon has a single tentacle (Figure 1A), which is known as a palpacle [5]. The palpacle is, in contrast to the gastrozooid tentacle, unbranched and nematocysts can be found along its entire length. As in gastrozooids, strong expression was detected for all marker-genes in young palpons within the growth zone, and expression disappeared from later developmental stages (Figures 2A–B). Expression was absent from mature palpons (Figures 2A, 4A, Additional files 4I, 5H, 6I), except for *nanos-2*, which remained expressed in a small domain at the proximal end of the palpon (Figure 4C). Unlike in gastrozooids, this *nanos-2* expression domain did not extend around the entire zooid but was restricted to a small patch close to the palpacle base (Figure 4C). Semi-thin sections indicated this patch as a site of nematogenesis (Figure 4D), suggesting that it is equivalent to the basigaster of gastrozooids. These similarities between gastrozooids and palpons were consistent with the hypothesis that palpons are derived gastrozooids that lost the ability to feed, i. e. they lack a mouth opening [5]. Expression of all marker-genes was found at the proximal end of the palpacle (Figure 4A-C, Additional files 2G, 3J, 4I, 5H, 6I, 6K). Densely packed, undifferentiated cells with i-cell characteristics were present within palpacle bases (Figure 4D). Additional secondary palpons are added at the anterior end of mature cormidia, and gonodendra form laterally of these secondary palpons [5]. We found small buds with marker-gene expression anteriorly from the youngest primary palpon, which were at the sites where these secondary structures arise (Figure 4E–G).

**Figure 4.**
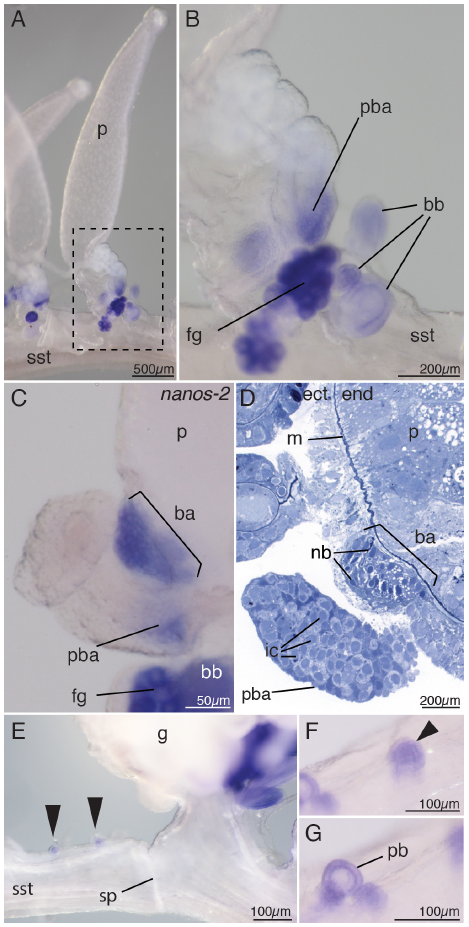
Palpons. (A-B) *vasa-1* transcript stained blue. (A) Mature palpon with *vasa-1* expression present exclusively in structures at the base. Anterior to the right. (B) Close up of the boxed region in A. *vasa-1* expression is restricted to the proximal end of the palpacle base, developing bracts, and young female gonophores. (C) *nanos-2* transcript in the basigaster region and the palpacle base. Anterior to the left. (D) Semi-thin longitudinal section of the palpon base, stained with toluidine blue, reveals interstitial cells in the palpacle base and developing nematocysts in the basigaster region. Anterior to the left. (E) Palpon buds at the anterior end of a cormidium (black arrows) express *vasa-1*. The sphincter region marks the posterior end of a cormidium. At the site of the sphincter the hollow stem can be constricted. Anterior to the right. (F) Close-up of an early palpon cluster bud (arrowhead). (G) Close-up of a later developmental stage of a palpon cluster with the palpon bud visible in the center and further buds laterally. ba: basigaster; bb: bract bud; ect: ectoderm; end: endoderm; fg: female gonodendron; ic: interstitial cell; m: mesoglea; p: palpon; pb: palpon bud; pba: palpacle base; sp: sphincter; sst: siphosomal stem.

Bracts are protective zooids, which can be found laterally along the siphosomal stem but also associated with palpons and gastrozooids (Figure 1B, [5, 7]). They are of scale-like morphology and function as protective shields. As in gastrozooids and palpons, marker-gene expression was found in early developing bract buds (Figure 4B) but was absent from older bracts once the typical bract morphology became obvious (Figure 2A).

### I-Cells and Germ Line Cells in Sexual Zooids

While some siphonophore species are dioecious, a colony of *Nanomia bijuga* produces gametes of both sexes [5]. Gametes are produced by gonophores, each of which is male or female. These gonophores are arranged into groups called gonodendra [5], which each exclusively bear male or female gonophores. Gonodendra are attached directly to the stem and develop laterally at the base of the palpon peduncle. There are gonodendra of both sexes associated with each palpon, one male and up to two female. The locations of these male and female gonodendra alternate between adjacent palpons (Figures 1A, 5A, [5]).

**Figure 5.**
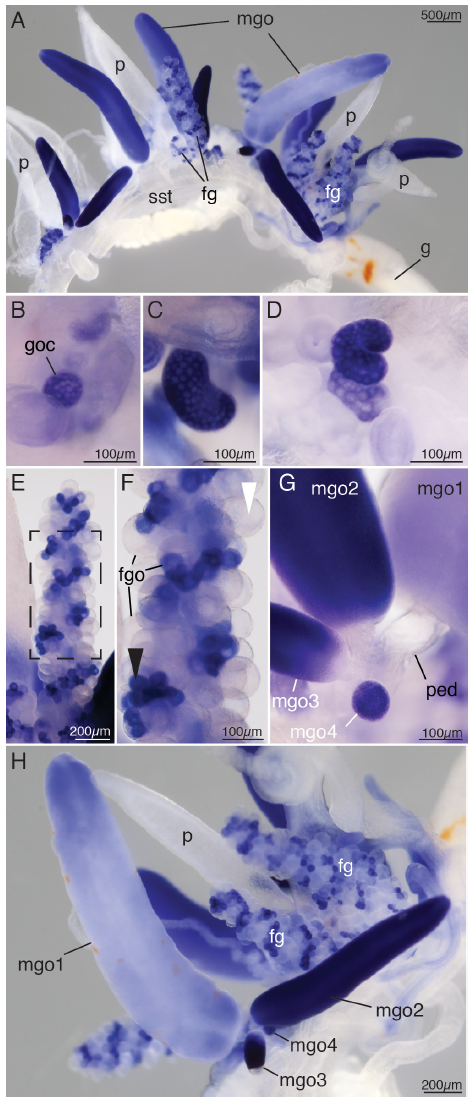
Gonodendra development, *vasa-1* transcript stained blue. (A) Mature cormidium, including male and female gonodendra. Anterior is to the left of the pane, ventral to the top of the pane. (B-E) Ontogenetic series of developing female gonodendra. (B) Cell cluster with *vasa-1* expression at the site of gonodendron formation at the base of a palpon. (C) Developing bean-shaped female gonodendron. (D) Developing female gonodendron starting to spiral. (E) Mature female gonodendron with developing gonophores with marker gene expression and mature gonophores with marker gene expression absent. (F) Close-up of female gonodendra (boxed area in E) with developing (black arrowhead) and mature gonophores (white arrowhead). Distal is up. (G) Close-up of the base of a male gonodendron. Later gonophores bud off the peduncle of the primary gonophore. The primary male gonophore (mgo1) is visible to the right. (H) Male gonodendron with an ontogenetic series of male gonophores, labeled mgo1-4 from oldest to youngest. *vasa-1* transcript abundance decreased as the male gonophore matured. fg: female gonodendron; fgo: female gonophore; g: gastrozooid; goc: gonodendron cell cluster; mgo: male gonophore; mgo1: oldest male gonophore; p: palpon; ped: peduncle; sst: siphosomal stem.

Female gonodendron formation has been described previously [27] as follows: Female gonodendra start to form as small buds protruding at the base of the palpon peduncle. Germ cells develop in between endoderm and ectoderm. Each gonophore within the female gonodendron contains a single egg. The egg is enclosed by a thin ectodermal layer within the developing female gonophore. Two lateral canals form from endodermal epithelial cells within the gonophore. The mature gonophore is attached to the blind-ending central stalk of the gonodendron by a delicate peduncle.

Close to the growth zone, the first indication of gonodendron development was round clusters of cells with strong marker-gene expression on the stem at the base of the young palpons (Figures 1B, 5B, Additional file 3M). These clusters were visible before bud formation became obvious and male and female clusters were morphologically indistinguishable from each other at this stage. *In situ* hybridization revealed expression of all five marker-genes in a helical pattern in the mature female gonodendron. This pattern corresponds to a previously unobserved helical morphological organization (Figure 5, Additional files 3G, 3M-P, 4K, 5J, 6M). The gonodendron buds started to twist early in development and a stronger signal was observed on the outer side of the developing stalk away from the axis of the helix (Figure 5C–D, Additional file 3N-O). This pattern persisted during the first turns until the gonodendron took on the appearance of a dense grape-like structure. At this stage all marker-genes were strongly expressed in all gonophores along the gonodendron and the helical organization was not apparent. Helical organization became obvious again in later ontogenetic stages (Figure 5E–F, Additional files 3G, 3P, 4K, 5J, 6M) when marker-gene expression decreased in mature gonophores (Figure 5F). The presence of signal in immature gonophores distributed in a helical pattern along the gonodendron indicated that new gonophores were produced along one side of the entire twisted stalk of the gonodendron. The chirality of the helices changed with the site of attachment. Gonodendra attached on the left side of a palpon showed a clock-wise directionality of turns.

The male gonodendron starts with the formation of a primary gonophore, which is cone shaped. Secondary and tertiary gonophores bud off the delicate peduncle of the primary gonophore (Figure 5G). The male gonophore is an elongated structure with a massive population of putative germ cells in the ectoderm. All marker-genes were strongly expressed in young and medium-sized gonophores but signal intensity was lower or absent in gonophores close to or at maturity (Figure 5H, Additional files 3Q, 4M, 5L, 6N). The absence of graded signals along the proximal-distal axis suggest that sperm maturation took place along the entire gonophore.

### Nectosomal Growth Zone has A Similar Structure as the Siphosomal Growth Zone

*Nanomia bijuga*, like most other siphonophore species, has a nectosomal growth zone (Figure 1C) near the anterior end that produces the swimming zooids, called nectophores, which propel the whole colony through the water [5]. All five genes were strongly expressed in the nectosomal growth zone at the tip of the horn, in nectophore buds, and in young developing nectophores (Figures 6A–D, Additional file 5A). *In situ* hybridization and histological sections indicated the presence of i-cells in the thickened region of the nectosomal stem, the horn of the growth zone and young nectophore buds (Figure 6B,D–E). In case of *vasa-1*, the transcript persisted longest along the ridges of the nectophores (Figure 6A). Older nectophores were free of marker-gene transcripts (Figure 6B–C). In contrast to the other marker-genes, *nanos-2* expression was restricted to the very youngest nectophore buds (Figure 6C). In addition, in the stem subtending the growth zone the transcript was detected on the nectosomal stem in a salt and pepper pattern (Figure 6C). Sections revealed developing nematocysts in this region of the stem (Figure 6F). Undifferentiated cells with interstitial cell morphology were identified in the ectoderm of developing nectophores (Figure 6F–G).

**Figure 6.**
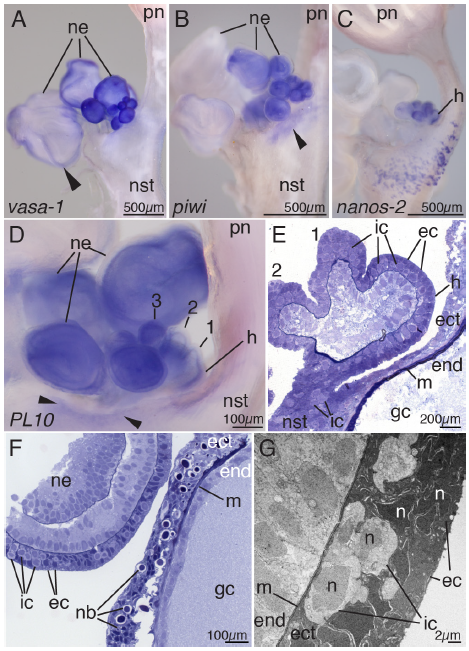
Nectosomal growth zone. Anterior is up in all figures. (A) *vasa-1* transcript. Transcription was longest detectable along the nectophore ridges (arrowhead). (B) *piwi* transcript. Marker gene expression was observed within the protruding nectosomal bulge (arrowhead), young buds, and developing nectophores. (C) *nanos-2* expression in the nectosomal horn and young developing buds. Signal on the nectosomal stem indicates sites of nematogenesis. (D) *PL10* transcript present in the nectosomal horn, youngest buds (1-3), young developing nectophores and within the protruding nectosomal bulge (arrowheads). Marker gene expression within the horn and young buds appeared strongest in deeper layers. (E) Semi-thin longitudinal section of early nectophore buds and the horn, stained with toluidine blue. Interstitial cells could be identified in the protruding bulge of the nectosomal stem, the horn and young developing buds (1-2). (F) Semi-thin longitudinal section in the region of the nectosomal horn showing nematogenesis in the ectoderm of the nectosomal stem subtending the growth zone and interstitial stem cells in the ectoderm of a developing young nectophore. (G) Transmission electron micrograph showing interstitial stem cells in in the interstices of the epithelial muscle cells within the ectoderm of a young nectophore. ec: epithelial cell; ect: ectoderm; end: endoderm; gc; gastric cavity; h: horn of the growth zone; ic: interstitial cell; m: mesoglea; n: nucleus; nb: nematoblasts; ne: nectophore; nst: nectosomal stem; pn: pneumatophore.

### Stem Cell Gene Expression is Found in A Subset of Regions with High Rates of Cell Proliferation

A qualitative assessment of cell proliferation revealed high densities of EdU labeled nuclei in domains with marker gene expression (compare Figure 2A and Figure 7A). Specifically, EdU labeled nuclei were found in the horns of both growth zones as well as in young buds and developing zooids both within the growth zones and along the siphosomal stem (Figure 7A–F). In all analyzed tissue samples the palpacle bases were consistently strongly EdU labeled in developing as well as in mature palpons (Figure 7F). Tentacle bases and developing tentilla were also strongly EdU labeled in gastrozooids (Figure 7A, 7G). In addition, high densities of EdU labeled nuclei were found in stem regions at the level and adjacent to both growth zones (Figure 7C,D,I), where only a few or no EdU labeled nuclei were identified in posterior regions of the nectosomal (Figure 7H) and siphosomal (Figure 7J) stem. These EdU labeled regions are the main sites of stem elongation in *Nanomia bijuga*. Interestingly, these stem regions were devoid of marker gene expression (compare Figures 2B and 7C, Figures 6A,B and 7D). Conspicuous cell division was occasionally observed along the dorsal midline of the stem (Figure 7K) whereas no signal was obtained in these regions in the *in situ* hybridizations. The number of EdU labeled cells in a particular zooid type decreased with level of maturity and in many cases no proliferative activity was found in mature zooids (Figure 7L–T). In developing male gonophores, our EdU assay showed a large number of dividing cells in the ectoderm (Figure 7 U–V). In developing female gonodendra, EdU labeled nuclei were consistently detected in developing gonophores bells (Figure 7W).

**Figure 7.**
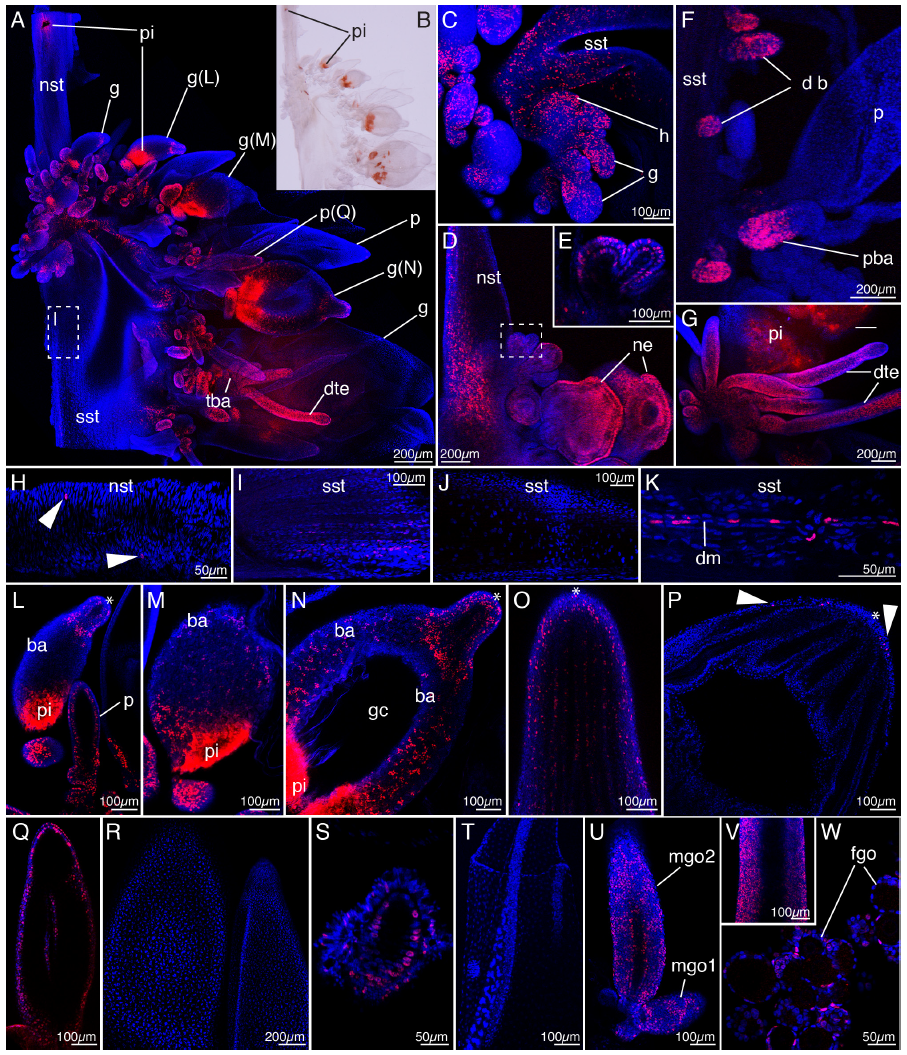
Assessment of cell proliferation in a mature colony after a five hour EdU pulse. Cells that divide during this interval incorporate EdU and their nucleii are stained red. All other nucleii appear blue. (A) Siphosomal growth zone. Anterior is up, ventral to the right. (B) Bright field image of tissue shown in A. Orange pigment spots are visible at the base of gastrozooids causing a red fluorescent signal visible in corresponding sites in A. (C) Close-up of the siphosomal growth zone shown in A showing EdU labeled nuclei in the horn, young developing buds and stem tissue. Anterior is up, ventral to the right. (D) Nectosomal growth zone with high densities of EdU labeled nuclei along the nectosomal stem, in young zooid buds as well as in developing nectophores. Anterior is up, dorsal to the right. (E) Youngest buds within the nectosomal growth zone, close-up of box in D. (F) Siphosomal stem fragment (SF2) with young developing buds, mature palpon and palpacle base. Anterior is up, ventral to the right. (G) Tentacle base and developing tentilla at the base of the gastrozooid basigaster. (H) Posterior part of the nectosomal stem of the colony shown in D. EdU labeled nuclei are sparsely scattered (arrowheads). Anterior to the left, lateral view. (I) EdU labeled nuclei along the stem posterior to the siphosomal growth zone, close-up of box in A. Anterior to the left, lateral view. (J) Siphosomal stem fragment (SF2) with EdU labeled nuclei absent. Anterior to the left, lateral view. (K) Siphosomal stem fragment (SF1) with EdU labeled nuclei along the dorsal canal. Anterior to the left, dorsal view. (L-P) Ontogenetic series of gastrozooid development. Apical is up. (L-N) Close-ups of gastrozooids g(L-N) shown in A. (O) Developing hypostome of a gastrozooid with high densities of EdU labeled nuclei. (P) Hypostome of a mature gastrozooid from siphosomal fragment (SF2) with few EdU labeled nuclei (arrowheads). (Q) Close-up of developing palpon p(Q) shown in A. Apical is up. (R) Two mature palpons from siphosomal fragment SF2 with EdU labeled nuclei absent. Apical is up. (S) Young developing bract from siphosomal fragment (SF2) with EdU labeled nuclei. Apical is up. (T) Mature bract from siphosomal fragment (SF3) with EdU labeled nuclei absent. Apical is up. (U) Developing gonophores from siphosomal fragment SF2. Apical is up. (V) Mid-section of a late male gonophore from siphosomal fragment SF3. Apical is up. (W) Developing female gonophores. Apical is up. b: bract; ba: basigaster; db: developing bud; dte: developing tentilla; dm: dorsal midline; fgo: female gonophore; g: gastrozooid; gc: gastric cavity; h: horn of the growth zone; mgo: male gonophore; ne: nectophore; nst: nectosomal stem; p: palpon; pba: palpacle base; pi: pigment; sst: siphosomal stem; *: apical end of gastrozooid; magenta: nuclei of cells, which formed during a five hour interval; blue(DAPI): nuclei, which did not divide during a five hour interval.

## Discussion

### A Cellular Perspective on Differences in Growth and form Between Siphonophores and Other Hydrozoans

The distribution of cell proliferation and i-cells in siphonophores differs in several key respects from other hydrozoans. These differences may play a critical role in the unique colony-level development and morphology of siphonophores.

Cell proliferation is high within the horn and young developing zooids of the siphonophore *Nanomia bijuga*. It then decreases in the course of zooid development and was mostly absent in mature zooids. This is in contrast to benthic colonial relatives such as *Tubularia* and *Hydractinia* where mitotic activity is maintained in mature gastrozooids [28]. Cell proliferation persists at the base of tentacles and palpacles, indicating continuous growth, as well as in gamete producing zooids. We also see rates of high proliferation in the stem within the growth zones. This is the first confirmation of restricted growth in the stem.

I-cells are found in a subset of the regions with high rates of cell proliferation. These regions include the horn and developing zooids (summarized in Figure 8). Interestingly, we do not detect i-cells along the stem of the colony, either within the growth zone or in the stem between mature zooids. This has several important implications. First, it suggests that epithelial cell division is responsible for stem elongation in *N. bijuga*, and does not require i-cells. Second, the lack of i-cells in the stem differs from benthic colonial hydrozoans that have widely distributed i-cells along their stolons and plastic growth. In the hydrozoan *Hydractinia echinata* i-cells reside in the stolon system which interconnects the different bodies of the colony [2, 10, 11]. These i-cell distribution patterns allow for addition of new zooids at various sites along the entire stolon system and different colonies of the same species do not have the exact same organization of zooids relative to each other.

**Figure 8.**
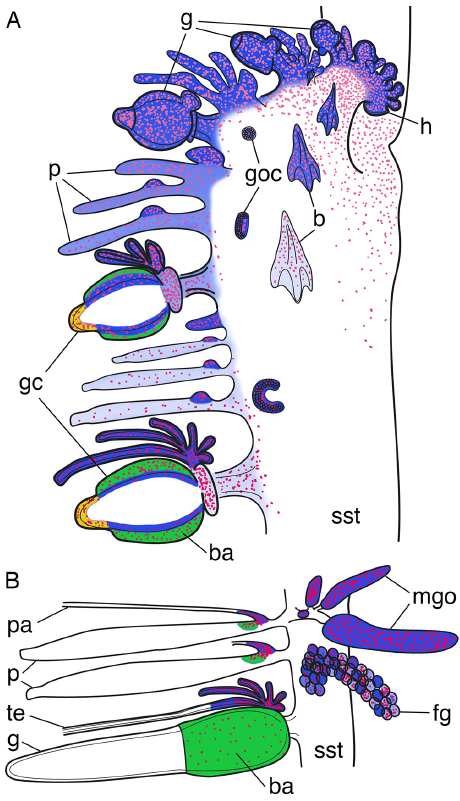
Schematic of i-cell distribution and cell proliferation in *Nanomia bijuga*. Shades of blue indicate density of i-cells, as indicated by coexpression of all marker genes. Magenta dots indicate density of cell proliferation. (A) Siphosomal growth zone and anterior part of the siphosome. Two gastrozooids (gc) are represented as cross sections. I-cells (blue) get restricted to deeper layers of the basigaster. Two marker genes continue to be expressed at locations in addition to those where all the marker genes are expressed. *nanos2* (green) is expressed in outer layers of the basigaster. *vasa-1* (yellow) is expressed within the developing hypostome. (B) Older siphosomal stem fragment with a mature gastrozooid, two palpons and developing gonozooids. Cell proliferation is spatially restricted to tentacle and palpacle bases and to developing gonophores. b: bract; fg: female gonodendron; g: gastrozooid; gc: gastrozooid cross section; goc: gonodendral i-cell cluster; h: horn; mgo: male gonophore; p: palpon; pa: palpacle, sst: siphosomal stem; t: tentacle.

The elongating stem of the siphonophore colony corresponds to the proximal part of the primary polyp, which is formed early in development (Figure 1D, [5]). Our data indicate that pluripotent i-cell populations get restricted to the sites of future growth zones during development of the primary polyp in the course of growth zone establishment. The fact that no marker gene expression can be found along the stem or in mature zooids bodies in combination with ceased mitotic activity in these older tissues in the colony may explain previous observations of reduced regenerative capacities in siphonophores relative to other colonial animals [29]. The restriction of stem cell pools to budding zones stands out as the key innovation, which lead to a reduced plasticity, enabled a far more precise, unique and highly organized model of growth observable in siphonophores.

### The Origin, Fate, and Developmental Potential of Siphonophore i-Cells

The potency, i.e. the ability to differentiate into other cell types, of siphonophore i-cells remains unknown. The interstitial cell lineage in many hydrozoans comprises pluripotent i-cells that give rise to unipotent progenitor cells [9, 30]. These stem cells give rise to somatic cells such as nerve cells, gland cells, nematocytes or gametes [2, 30, 31]. Previous work has revealed diversity within Hydrozoa in the potency of i-cells. In the freshwater polyp *Hydra*, i-cells are pluripotent but cannot give rise to epithelial cells [31–33]. In contrast, the i-cells of the marine colonial hydrozoan *Hydractinia* can give rise to all cell types including epithelial cells [2, 10, 11]. Despite the fact that we do not provide double labelling experiments, we interpret the sites, which show expression for all marker genes as the sites where pluripotent i-cells reside (summarized in Figure 8). In several cases we identified differentials in the expression domains of different markers, which we interpret as hints for the presence of determined progenitor cells. For instance, *vasa1* continues to be expressed in the tip of the developing gastrozooid (Figure 8) and the expressing cells may give rise to a tissue specific yet unidentified cell type. Exclusive expression of *nanos2* indicates the presence of nematoblast since a role of *nanos2* in nematocyst formation pathways has been demonstrated previously (Figure 8) [23]. Migrating progenitor cells may be the mechanism by which the elongating stem gets replenished with somatic stem cells such as nerve cells, since these cell types are present within stem tissue [34–36]. Some of the mitotically active cells in the stem region may therefore be amplifying progenitor cells which have lost i-cell specific signatures. Two giant nerve fibres run along the dorsal midline of the siphosomal stem of *N. bijuga*, which function as rapid conduction pathways and for which a syncytial character has been reported [34, 37]. Dividing cells along this dorsal midline (Figure 7K) in older parts of the colony may point to the presence of nerve progenitor cells ultimately contributing to giant axon fibres. The cellular identity of these cells could, however, not be established in this study. Migratory activity of progenitor cells with already determined fates, e.g. nematoblasts and neuroblasts, has been frequently demonstrated in of a variety of hydrozoan species (e.g. [38–42].

This study did not differentiate between several possible origins for the i-cells that give rise to additional secondary palpons and gonodendra at the anterior end of each cormidium (Figure 4E–G). It could well be that these cells are incorporated into the developing cormidium early in the growth zone. Alternative explanations could involve dedifferentiation, where differentiated cells revert back to an undifferentiated i-cell, or migration of i-cells. Migration, however, would require marker gene expression along the stem of the colony for which we did not find evidence.

### The Similarity of Siphonophore Growth Zones to Plant Meristems

Others have likened regions of growth in hydrozoans to land plant meristems and suggested localized growth at tips of stolons or hypostomes [43, 44]. Such localized cell proliferation and meristematic character of these regions could, however, not be confirmed in later studies [45, 46]. Cell proliferation was rather found present along the entire stolon systems [45–48]. Berking et al. (2002) described a meristem-like organ in the thecate Hydrozoan *Dynamena pumila*. In a *D. pumila* colony each stem has a growing tip, which, usually, neither ends in a stolon tip nor as a polyp but grows out as to form the stem [49]. This growing tip frequently splits into three primordia, two of which give rise to lateral buds, which develop into polyps, and the third forming a new growing stem tip.

Within Hydrozoa, siphonophores seem to have taken the degree of spatial restriction of a pluripotent pool of cells and proliferating cells to an extreme. This makes the plant analogy a particularly interesting one though the cellular organization in plants and hydrozoans clearly differs. Analogously to a plant meristem, which produces structures that develop into functional organs, siphonophore buds generated laterally from the horn within the growth zone develop into specialized bodies. Both the meristem in plants and the horns within the growth zones can be characterized as restricted morphogenetic fields that harbor constantly dividing cells. In both cases new cells are produced for expansion and tissue differentiation. In the case of the siphonophore horn, cell division of endodermal and ectodermal cells as well as nested amplifying interstitial cells generate tissue available for bud formation. Cells differentiate and zooids mature as these newly formed structures are carried away from the horn. Our observations of cellular proliferation and stem cell distribution within the siphonophore colony allow for a more detailed extension of the analogy from observable patterns to the cellular dynamics that give rise to those patterns.

## Conclusions

Siphonophores may have realized a complex and precise colony-level development and organization by restricting the sites of zooid formation and differentiation through the spatial restriction of interstitial stem cells. In siphonophores restriction of growth to particular zones might have led to precise growth and a loss of plasticity through the restriction of developmental and regenerative capacities in older parts of the colony.

## List of Abbreviations

i-cell: interstitial stem cell, EdU: 5-ethynyl-2-deoxyuridine, FSW: filtered sea water, ROV: remotely operated vehicle, TEM: transmission electron microscopy

## Authors’ Contributions

SS and CD designed the study. SS, FG and SH collected specimens. SS, FG and PB performed *in situ* hybridizations. SC and SS performed histological analysis. SS performed the cell proliferation assay. FZ conducted phylogenetic analyses. SS, FG, SC, PB, FZ, SH and CD analyzed the data. SS and CD wrote the paper. All authors have read, revised and approved the final manuscript.

## Acknowledgments

We thank Leo W. Buss and Uri Frank for critical feedback on a preprint version of the manuscript. SS and CWD thank Claudia Mills for informing us about high abundances of *Nanomia bijuga* at Friday Harbor Labs (FHL), San Juan Island, WA, and Billie Swalla for hosting SS at her lab. We thank members of the Dunn lab for discussion and feedback on the manuscript. We also thank the MBARI crews and ROV pilots for collection of *N. bijuga* specimens. Computational work was conducted at the Center for Computation and Visualization, Brown University. This research was supported by the US National Science Foundation (grant 1256695 and the Alan T. Waterman Award) and by the David and Lucile Packard Foundation.

## References

1. DunnC: Siphonophores. Curr Biol 2009, 19:R233–R234.

2. PlickertG, FrankU, MüllerWA: Hydractinia, a pioneering model for stem cell biology and reprogramming somatic cells to pluripotency. Int J Dev Biol 2012, 56:519–534.

3. Harvell CD: The evolution of polymorphism in colonial invertebrates and social insects. Q Rev Biol 1994, 96:155–185.

4. BoardmanRS, CheethamAH: Degrees of colony dominance in stenolaemate and gymnolaemate Bryozoa. In Animal Colonies: Development and Function through Time. Volume 603. Edited by BoardmanRS, CheethamAH, OliverWA; 1973:121–220.

5. Totton AK: A Synopsis of the Siphonophora. London: British Museum (Natural History); 1965.

6. SiebertS, PughPR, HaddockSHD, DunnCW: Re-evaluation of characters in Apolemiidae (Siphonophora), with description of two new species from Monterey Bay, California. Zootaxa 2013, 3702:201–232.

7. DunnCW, WagnerGP: The evolution of colony-level development in the Siphonophora (Cnidaria:Hydrozoa). Dev Genes Evol 2006, 216:743–754.

8. Dunn CW: Complex colony-level organization of the deep-sea siphonophore Bargmannia elongata(Cnidaria, Hydrozoa) is directionally asymmetric and arises by the subdivision of pro-buds. Dev Dyn 2005, 234:835–845.

9. WeismannA: The origin of the sexual cells in hydromedusae (Foreign title: Die Entstehung der Sexualzellen bei Hydromedusen). Gustav Fischer 1883:1–422.

10. MüllerWA, TeoR, FrankU: Totipotent migratory stem cells in a hydroid. Dev Biol 2004, 275:215–224.

11. KünzelT, HeiermannR, FrankU, MüllerW, TilmannW, BauseM, NonnA, HellingM, SchwarzRS, PlickertG: Migration and differentiation potential of stem cells in the cnidarian Hydractinia analysed in eGFP-transgenic animals and chimeras. Dev Biol 2010, 348:120–129.

12. LeclèreL, JagerM, BarreauC, ChangP, Le GuyaderH, ManuelM, HoulistonE: Maternally localized germ plasm mRNAs and germ cell/stem cell formation in the cnidarian Clytia. Dev Biol 2012, 364:236–248.

13. Lentz TL: The fine structure of differentiating interstitial cells in Hydra. Z Zellforsch 1965, 67:547–560.

14. MochizukiK, SanoH, KobayashiS, Nishimiya-FujisawaC, FujisawaT: Expression and evolutionary conservation of nanos-related genes in Hydra. Dev Genes Evol 2000, 210:591–602.

15. SeipelK, YanzeN, SchmidV: The germ line and somatic stem cell gene Cniwi in the jellyfish Podocoryne carnea. Int J Dev Biol 2004, 48:1–7.

16. RebscherN, VolkC, TeoR, PlickertG: The germ plasm component vasa allows tracing of the interstitial stem cells in the cnidarian Hydractinia echinata. Dev Dyn 2008, 237:1736–1745.

17. DunnCW, HowisonM, ZapataF: Agalma: an automated phylogenomics workflow. BMC Bioinformatics 2013, 14:330.

18. KernerP, DegnanSM, MarchandL, DegnanBM, VervoortM: Evolution of RNA-Binding Proteins in Animals: Insights from Genome-Wide Analysis in the Sponge Amphimedon queenslandica. Mol Biol Evol 2011, 28:2289–2303.

19. Edgar RC: MUSCLE: multiple sequence alignment with high accuracy and high throughput. Nucleic Acids Res 2004, 32:1792–1797.

20. StamatakisA: RAxML-VI-HPC: maximum likelihood-based phylogenetic analyses with thousands of taxa and mixed models. Bioinformatics 2006, 22:2688–2690.

21. FelsensteinJ: Cases in which parsimony or compatibility methods will be positively misleading. Syst Zool 1978, 27:401–440.

22. GenikhovichG, KurnU, HemmrichG, BoschTCG: Discovery of genes expressed in Hydra embryogenesis. Dev Biol 2006, 289:466–481.

23. KanskaJ, FrankU: New roles for Nanos in neural cell fate determination revealed by studies in a cnidarian. Journal of cell Science, 126:3192–3203.

24. SiebertS, RobinsonMD, TintoriSC, GoetzF, HelmRR, SmithSA, ShanerN, HaddockSHD, DunnCW: Differential Gene Expression in the Siphonophore Nanomia bijuga (Cnidaria) Assessed with Multiple Next-Generation Sequencing Workflows. PLoS ONE 2011, 6:e22953.

25. OzbekS, PokidyshevaE, SchwagerM, SchulthessT, TariqN, BarthD, MilbradtAG, MoroderL, EngelJ, HolsteinTW: The Glycoprotein NOWA and Minicollagens Are Part of a Disulfidelinked Polymer That Forms the Cnidarian Nematocyst Wall. J Biol Chem 2004, 279:52016–52023.

26. MackieGO, PughPR, PurcellJE: Siphonophore Biology. Adv Mar Biol 1987, 24:97–262.

27. CarréD: Etude histologique du developpement de Nanomia bijuga (Chiaje, 1841), siphonophore physonecte, Agalmidae. Cah Biol Mar 1969, 10:325–341.

28. Campbell RD: Cell proliferation and morphological patterns in the hydroids Tubularia and Hydractinia. J Embryol exp Morph 1967, 17:607–616.

29. MackieGO, BoagDA: Fishing, Feeding and Digestion in Siphonophores. Pubbl statz zool Napoli 1963, 33:178–196.

30. Bode HR: The interstitial cell lineage of hydra: a stem cell system that arose early in evolution. Journal of cell Science 1996, 109:1155–1164.

31. BoschTCG, DavidCN: Stem cells of Hydra magnipapillata can differentiate into somatic cells and germ line cells. Dev Biol 1987, 121:182–191.

32. CampbellRD, DavidCN: Cell cycle kinetics and development of Hydra attenuata. II. Interstitial cells. Journal of cell Science 1974, 16:349–358.

33. Bosch TCG: Hydra and the evolution of stem cells. BioEssays 2009, 31:478–486.

34. Mackie GO: Report on giant nerve fibres in Nanomia. Publications of the Seto Marine Biological Laboratory 1973:745–756.

35. GrimmelikhuijzenCJP, SpencerAN, CarreD: Organization of the nervous system of physonectid siphonophores. Cell Tissue Res 1986, 246:463–479.

36. ChurchSH, SiebertS, BhattacharyyaP, DunnCW: The Histology of Nanomia bijuga (Hydrozoa: Siphonophora). bioRxiv 2014. doi:10.1101-010868.

37. Mackie GO: Coordination in Physonectid Siphonophores. Mar Behav Physiol 1978, 5:325–346.

38. HagerG, DavidCN: Pattern of differentiated nerve cells in hydra is determined by precursor migration. Development 1997, 124:569–576.

39. DenkerE, ManuelM, LeclereL, Le GuyaderH, RabetN: Ordered progression of nematogenesis from stem cells through differentiation stages in the tentacle bulb of Clytia hemisphaerica (Hydrozoa, Cnidaria). Dev Biol 2008, 315:99–113.

40. FujisawaT: Role of interstitial cell migration in generating position-dependent patterns of nerve cell differentiation in hydra. Dev Biol 1989, 133:77–82.

41. BoehmA-M, BoschTCG: Migration of multipotent interstitial stem cells in Hydra. Zoology 2012, 115:1–8.

42. HeimfeldS, BodeHR: Interstitial Cell Migration in Hydra attenuata II. Selective Migration of Nerve Cell Precursors as the Basis for Position-Dependent Nerve Cell Differentiation. Dev Biol 1984, 105:10–17.

43. Bonner JT:Morphogenesis. Princeton, N.J.: Princeton University Press; 1952:vii–296.

44. Berrill NJ: The Polymorphic Transformations of Obelia. Quarterly J Microsc Sci 1949, 90:235–264.

45. Kosevich IA: Morphogenetic foundations for increased evolutionary complexity in the organization of thecate hydroids shoots (Cnidaria, Hydroidomedusa, Leptomedusae). Biol Bull Russ Acad Sci 2012, 39:172–185.

46. BravermanM: Studies on hydroid differentiation VII. The hydrozoan stolon. J Morphol 1971, 135:131–152.

47. HaleLJ: CellMovements, Cell Division and Growth in the Hydroid Clytia johnstoni. J Embryol Exp Morphol 1964, 12:517–579.

48. Suddith RL: Cell Proliferation in the Terminal Regions of the Internodes and Stolons of the Colonial Hydroid. Campanularia flexuosa. Amer Zool 1974, 14:745–755.

49. BerkingS, HesseM, KH: A shoot meristem-like organ in animals; monopodial and sympodial growth in Hydrozoa. Int J Dev Biol 2002, 46:301–308.

50. HaddockSHD, DunnCW, PughPR: A re-examination of siphonophore terminology and morphology, applied to the description of two new prayine species with remarkable bio-optical properties. J Mar Biol Ass U K 2005, 85:695–707.

51. Nanomia bijuga whole animal and growth zones. http://commons.wikimedia.org/wiki/File:Nanomia_bijuga_whole_animal_and_growth_zones.svg. Accessed 2013-11-01.

52. Nanomia life cycle. http://commons.wikimedia.org/wiki/File:Nanomia_life_cycle_vector_wikimedia.svg. Accessed 2013-11-01.

